# Computer vision guided open-source active commutator for neural imaging in freely behaving animals

**DOI:** 10.1101/2024.05.28.596351

**Authors:** Ibrahim Oladepo, Kapil Saxena, Daniel Surinach, Malachi Lehman, Suhasa B. Kodandaramaiah

## Abstract

Recently developed miniaturized neural recording devices that can monitor and perturb neural activity in freely behaving animals have significantly expanded our knowledge neural underpinning of complex behaviors. Most miniaturized neural interfaces require a wired connection for external power and data acquisition systems. The wires are required to be commutated through a slip ring to accommodate for twisting of the wire or tether and alleviate torsional stresses. The increased trend towards long term continuous neural recordings have spurred efforts to realize active commutators that can sense the torsional stress and actively rotation the slip ring to alleviate torsional stresses. Current solutions however require addition of sensing modules. Here we report on an active translating commutator that uses computer vision (CV) algorithms on behavioral imaging videos captured during the experiment to track the animal’s position and heading direction in real-time and uses this information to control the translation and rotation of a slipring commutator to accommodate for accumulated mouse heading orientation changes and position. The CV guided active commutator has been extensively tested in three separate behavioral contexts and we show reliable cortex-wide imaging in a mouse in an open-field with a miniaturized widefield cortical imaging device. Active commutation resulted in no changes to measured neurophysiological signals. The active commutator is fully open source, can be assembled using readily available off-the-shelf components, and is compatible with a wide variety of miniaturized neurophotonic and neurophysiology devices.

## 1.0 Introduction

Understanding how the brain mediates complex behaviors requires the synchronized acquisition of both large-scale neural activity and behavior monitoring. Miniaturized neural devices that can be docked to mice, for neural recording and active manipulation of activity have become key tools for neuroscience to address these questions.

Miniaturized neurophotonic imaging devices allow imaging of specific populations of neurons at large scale have been particularly instrumental in conducting these critical neuroscience studies^1–3^. The generally open-source culture underpinning these innovations has spurred the development of myriad miniaturized neurophotonic with new variants of miniscopes that have smaller form factors and multiple FOVs^4,5^, large FOVs^1,6^. Variants that incorporate optics for structured optogenetic stimulation^7^, and simultaneous electrophysiology recording^8^ have also been developed. These tools complement the head borne electrophysiology recording devices such as tetrode Microdrive^9–14^, and more recent miniaturized high-density CMOS recording probes^15–19^.

Most miniaturized head borne devices require a wired tether for powering the devices, and interfacing with a fixed remote data acquisition system. The wire can get entangled as behaving mice locomote naturally in the behavior arenas, or when they perform tasks requiring them to execute several stereotyped behavioral trajectories (such as in a 8-maze choice task^20^). This is typically mitigated by using passive commutators that can relieve the torsion in the wires and transferring data through a slip ring. The increased trend towards long term continuous neural recordings have spurred efforts to realize active commutators. These active commutators rely on inertial.

Both these solutions require incorporation of additional hardware elements within the head borne devices which constrains the design space where functional elements of the head borne device needs to be limited to < 3g (∼15% of bodyweight of a 20g mouse). Wireless data transfer rates are typically limited, and further require specialized hardware surrounding the behavioral arena for powering and data transfer.

To mitigate this issue, active commutator systems utilizing inertial measurement unit (IMU)^21^, magnetic rotation sensor^22^, torque sensor^14^, hall sensor^23^, and video-based tracking^24^ have been developed. Measurement of an animal’s heading angle by using a torque sensor to estimate animal rotations (by indirectly measuring tethered cable rotations) is easy to implement, but the effectiveness of this approach is limited by sensitivity issues, as cables are poor transmitters of torque^14,21^. Similarly, hall sensor and magnetic rotation sensor-based heading angle estimation approaches are affected by sensitivity. The IMU-based heading angle estimation approach is not affected by the cable sensitivity issue, but requires calibration due to magnetic distortions^21^. Further, the IMU data can necessitate adding more channels to the commutator for data transfer, adding more weight to the already limited payload and limited cable data transfer bandwidth.

Video-based methods offer potential solutions to issues with torque, or inertial sensing and has been explored previously where LEDs on the head-mounted devices could track the position and direction of a mouse in the captured video frames^24^. The incorporation of LEDs however requires modification to existing miniature imaging and recording technologies. Given that monitoring the behavior of the animal is a critical requirement, and in these experiments, making use of this information to actively rotate a commutator and further move the commutator along with a mouse across a large behavior arena could enable new kinds of experiments with existing miniaturized devices.

Here we present a computer vision (CV) guided active translating commutator. The CV guided active commutator leverages recent advances in real-time markerless tracking to estimate the location and heading direction of the animal and uses a translating stage and a motor to move and rotate a slip ring commutator along with the mouse. We show that position and heading direction can be computed in real-time at 6 Hz in three different behavioral assays - open field behavior, active place avoidance behavior and the Barnes maze spatial navigation task. We further show that this information can be used to actively control the position and angular orientation of the slip ring commutator in response to mouse locomotion in the APA task and exploration of a linear maze.

## 2.0 Methods

### 2.1 General Design of the CV-guided active commutator

The CV-guided active commutator system dynamically adjusts the position and angular orientation of a slip ring commutator in response to estimated movement and angular orientation of a mouse within a behavioral arena. The overall principle of operation is illustrated in **Figure 1A**. An overhead camera captures video of the mouse behaving in the arena. A deep neural net algorithm (DeepLabCut^25^) is used to estimate the real-time position and heading angle direction of the mouse. These estimates of the mouse position and angular heading direction are used to actively translate and rotate a slip ring commutator.

**Figure 1.**
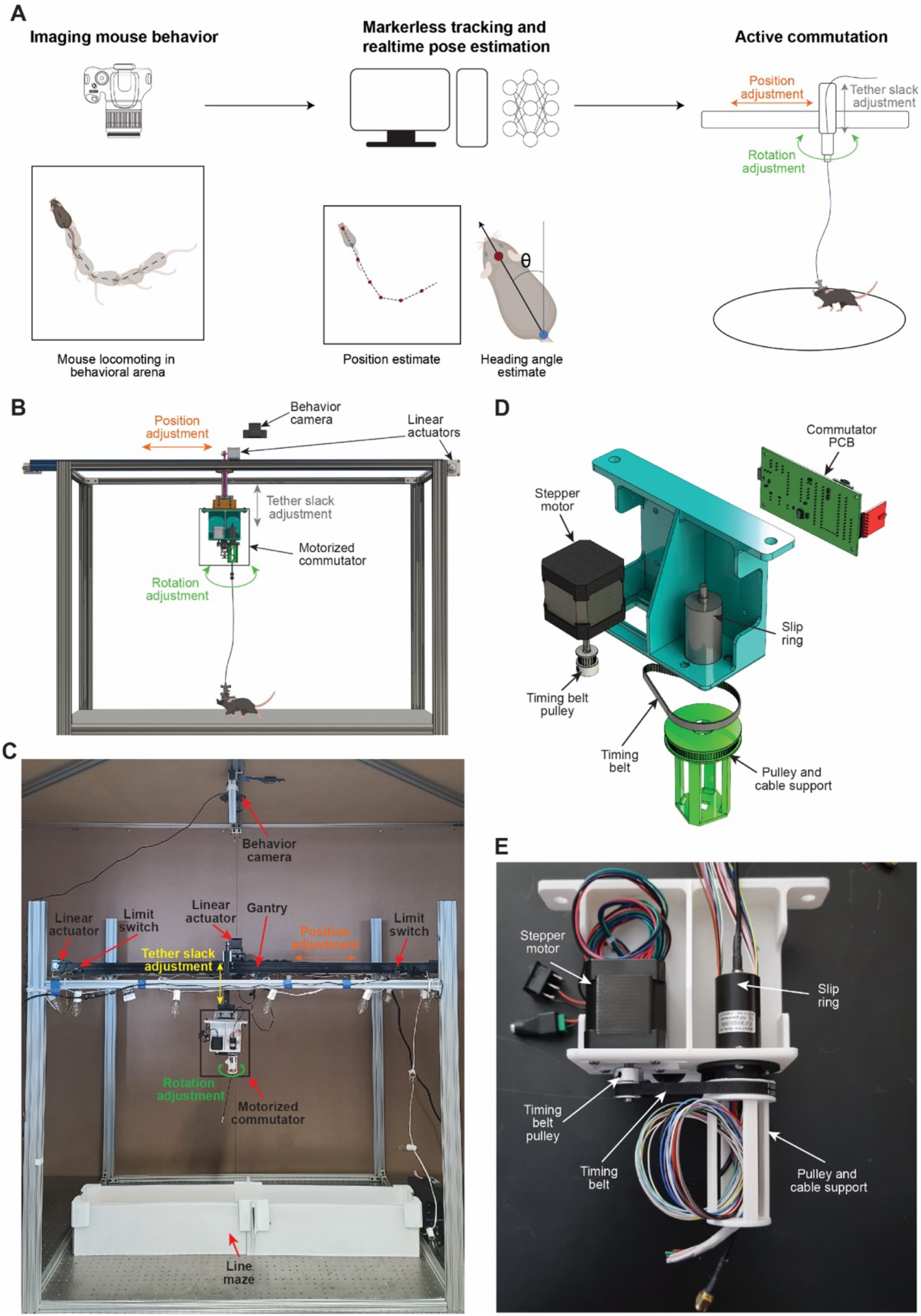
An open-source CV-guided active, translating commutator system: (A) Principle of operation - mouse behavior video captured from an overhead camera is used to estimate real-time position of the mouse location and heading direction, which is used to actively translate and rotate a slip ring commutator. (B) A computer aided design (CAD) schematic of the CV-guided active translating commutator system. (C) Photo of the CV-guided active translating commutator system. (D) Detailed CAD schematic of the motorized commutator module. (E) Photograph of the motorized commutator module.

The overall architecture of the CV-guided active commutator hardware is shown in **Figure 1B**. The CV-guided commutator consists of two main components, a behavior imaging camera located above an arena gantry that supports the actuator components. Two stepper motors (NEMA 17, OpenBuilds Parts Store) coupled to linear actuator modules provide actuation in x and z directions. A rotation module is mounted on the z-translation stage (**Figure 1B-C**). A third stepper motor (NEMA 17) is coupled to the slipring commutator via a timing belt drive (**Figure 1D**). We tested two slip rings: a single channel coax slip ring with 24 channel plane wires (LPC-24YT-2402-01HC, JINPAT Electronics) and a three-channel USB slip ring with 24 plane wires (LPT000-2402-04HF, JINPAT Electronics). The stepper motor and slip ring were attached to the anterior side of the 3D-printed holding structure. A custom printed circuit board (PCB) was designed to wirelessly receive commands from a computer and drive the stepper motor (**Figure S1**). The custom PCB housed a microcontroller (Teensy 4.0; **Figure S1**), a radio transceiver (NRF24L01, **Figure S1**) and a stepper motor driver (A4988, Pololu; **Figure S1**) and was mounted on the posterior side of the 3D-printed holding structure. **Figure 1D** shows a photograph of the rotation module of the CV-guided motorized commutator. The translation stages feature the same custom PCB used in the rotation stage and have stepper motors mounted directly on the arena gantry (**Figure 1B-C**). The x-translation stage is equipped with two limit switches on either end to prevent the mounted components from running past the edge and for self-calibration of the stage (**Figure 1C**).

### 2.2 Operating Modes

The CV-guided commutator can operate in two modes: passive and active. In the passive mode, the experimenter manually controls the commutator using a joystick and the commands are wirelessly transmitted to the commutator (**Figure S2**). In the passive mode, there is no computerized tracking of an animal. The control is solely based on video playback from cameras attached to the experiment arena. In the active mode, the commutator control is executed automatically by the computer without any input from the experimenter using location and heading angle estimates (**Figure S2**).

### 2.3 Training and evaluating real-time position and heading direction algorithms

The position and heading direction of the mice were estimated using the DeepLabCut (DLC) toolbox—an open-source pose estimation and behavioral analysis toolbox^25^. The toolbox was employed to train a model tailored to the arena’s environment that incorporates the CV-guided commutator. We trained three models, DLC implemented on MobileNetV2^26^, DLC implemented on ResNet^27^, and a third model that used SLEAP implemented on U-Net^28,29^. For comparing the real-time detection capabilities of these models, behavior videos from three different studies were used - mouse performing an active place avoidance task^30^ (n=3 mice), mice performing a spatial navigation task, Barnes maze^31^ (n=3 mice), and mice exploring an open field arena (n=3) mice. In each behavioral assay, 50 randomly selected frames from videos lasting 15 minutes were manually annotated to generate a ground truth of the location of the head and base of the tail. These were compared with the estimated head position and position of the base of the tail from the three models evaluated. In each assay, estimated and actual positions of head and tail base (manually annotated) were compared using a 10-pixel radius threshold. The estimation error was used as a metric for comparing the three models.

### 2.4 Implementation of rotation and translation compensation

To compensate for the rotation of the animal, the model estimates the heading angle. This was achieved by estimating the x and y coordinates of the head of the animal and the base of the tail of the animal across sequential camera frames captured at a rate of six to ten frames per second (FPS). The magnitude of frame-to-frame change in heading angle was computed using the dot product of the heading direction vectors, and the direction of the change, i.e., clockwise (CW) or counter-clockwise (CCW) was computed using the cross-product of the heading direction vectors. During operation, the cumulative change in heading direction was computed and when the cumulative value surpassed a predefined threshold, the rotation motor in the active commutator was activated to alleviate torsional stresses in the cable. We used a threshold of 90 degrees in the APA task and Barnes maze task, and 225 degrees in the linear maze task.

For translational compensation on the x stage, the model estimated the location of the head. Experiments were conducted when mice explored in a linear 1.2 m long maze. The length of the maze was virtually divided into eight segments, each ∼150 cm long. For each frame of the behavioral video, the estimate of the mouse’s head location was used to determine the current segment occupied by the mouse. When mice moved between segments, the linear stage was activated to ensure the commutator was directly above the mouse. Activation commutations were evaluated in the APA task and the linear maze.

### 2.5 Cortex-wide calcium imaging during active commutation

We performed cortex-wide mesoscale calcium imaging using a miniaturized microscope, mini-mScope^1^ through a polymer transparent cranial window^32,33^ during the APA task using the CV-guided active commutator to transmit signals through the slip ring. Calcium activities were acquired at 15 frames per second (FPS). The CMOS gain was set to a value of 55, and the LED voltage and current was set to 8V 0.8A for the blue LEDs. The blue LEDs were pulsed for 120 seconds, prior to the experiment, to allow them to warm up and reach a stable intensity. The mice were brought into the different behavioral arenas under red light and placed into an opaque cylinder at the center of the maze ∼90 seconds after the LEDs were turned on. The mini-mScope was attached to the mice via 3 interlocking magnets. At ∼120 seconds, the opaque cylinder was removed, marking the start of the trial.

### 2.6 Calcium data pre-processing

Calcium imaging was captured under both blue light illumination and green light illumination in alternate frames. The mean pixel intensity of each frame captured by the mini-mScope was calculated, and K-means clustering was used to classify each mean pixel intensity of the video and segregate the frame captured under blue light illumination (GCaMP signals) and green light illumination (reflectance signals, See Rynes et al.^1^). K-means clustering also enables identification and removal of outlier frames in the data due to large motion artifacts or irregularities in LED intensity (∼0.04% of all frames). The videos corresponding to both illumination wavelengths were then passed through a motion correction algorithm^34^.

The calcium data videos were compressed to 80% of their original size with a bilinear binning algorithm (2022b, MathWorks). One frame randomly selected in each trial was used to draw a mask around the imaged brain surface and exclude the background and superior sagittal sinus artery to reduce noise in the overall DF/F signal. For each mouse, the masks across all trials were averaged to generate a mouse-specific average cortex mask. The average mask was imposed across images acquired in all trials for a mouse so that the number of pixels used in each analysis remained consistent.

Each pixel within the mask was corrected for global illumination fluctuations using a correction algorithm that produces DF/F data^35^. The DF/F data was filtered using a zero-order phase Chebyshev band-pass filter with cutoff frequencies of 0.1 Hz and 5 Hz (2022b, MathWorks). The resulting data was then spatially filtered with a 7-pixel nearest-neighbor average using a custom MATLAB (2022b, MathWorks) script. The resulting DF/F time series for each pixel was then z-scored.

## 3.0 Results

### 3.1 Comparative analysis of pose estimation toolboxes and networks

Accurately tracking animal position and heading direction across diverse experimental assays and imaging conditions is crucial for a CV-guided active commutator to reliably mitigate wire entanglement. Several open source markerless pose estimation models^25,28^ have demonstrated capabilities in accurately tracking mice across a wide range of imaging setups and behavioral assays. To determine the model most suitable for integration with a CV-guided active commutator, we compared three models—DLC implemented in MobileNetV2^25,26^, DLC implemented in ResNet^25,27^, and SLEAP implemented on U-Net^28,29^—based on four criteria. First, the selected markerless pose estimation model should be able to track the position of the mouse in a wide range of imaging conditions and behavioral assays. Second, the model should be accurate and reliable such that it ensures minimal tracking errors. Third, the implementation of the model should allow for near real-time (>6 Hz) estimation of the animal’s position and heading angle, allowing up to 10 compensatory adjustments to the commutator position and angular orientation per minute. Lastly, the model should be executable on a regular desktop computer without necessitating an expensive graphic processing unit (GPU), ensuring that the system is accessible and cost-effective for widespread laboratory use.

To evaluate the efficacy of the three models, we tested markerless tracking capabilities on three independent behavioral assays that can be tracked using a single overhead camera. In the APA task, mice are introduced into a circular rotation arena and receive mild foot shocks when entering a designated sector^30,36^. As the arena rotates, mice must continuously move and adjust their position to avoid entering these shock zones^36^. The behavioral camera encompassed a circular area of 34.5 cm diameter and had a bed of parallel wires for delivering shock. The Barnes Maze task, a dry-land version of a spatial task for studying effects of aging on navigation in rats, is an avoidance task in which rodents learn to leave the center of a large circular platform (where they are exposed under bright, aversive lights) to find an escape hole among those at the edge of the platform^31^. Only one of the holes provides access to a nest-space. The Barnes maze allowed the evaluation of real-time markerless tracking of an animal exploring a large open environment (1 m diameter). Third, we used a standard open field arena, where mice explored the whole extent of the maze, exhibiting a variety of behaviors, including thigmotaxis, rearing, and grooming.

**Figure 2A-C** shows results of markerless tracking of the mouse head position in each of the three behavioral assays, as well as the range of head angle estimated. As expected, in the APA task, active avoidance resulted in higher occupancy in the non-shock areas of the maze, with polarized heading angles. Barnes maze tasks resulted in animals predominantly exploring the edge of the maze, with nearly uniformly distributed heading angles. Similarly, in open field experiments, animal position could be estimated without any large errors or jumps in position estimates. Animals had a tethered head borne device, and tracking was robust to variations in wired connection position within the FOV.

**Figure 2.**
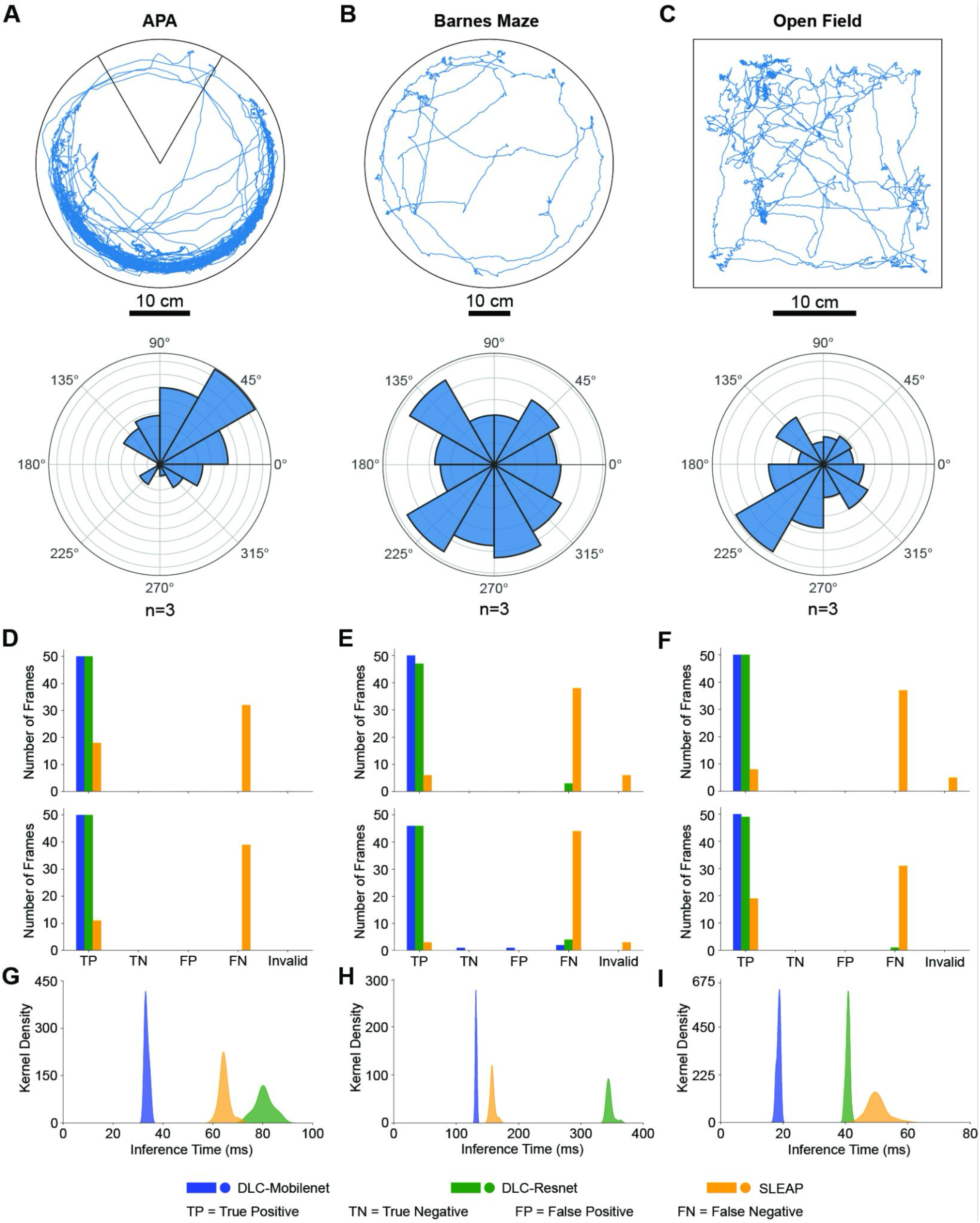
Evaluation of pose estimation toolboxes and convolution neural networks for realtime position and heading direction estimates. (A) Trajectory plot of a mouse performing the active place avoidance task^30^ (top) and polar histogram showing distribution of the head direction (n = 3 mice) in the same arena (bottom). The pose tracking was done using the DLC toolbox. (B) Trajectory plot of a mouse performing the Barnes maze task^31^ (top) and polar histogram showing distribution of the head direction (n = 3 mice) in the same arena (bottom). (C) Trajectory plot of a mouse in an open field arena (top) and polar histogram showing distribution of the heading direction (n = 3 mice) in the same (bottom). The pose tracking was done using the DeepLabCut toolbox. (D) Accuracy of real-time estimation of head position (top) and tailbase position (bottom) of the mouse during APA behavior for each of the three models: DLC implemented on MobileNetV2, DLC implemented on ResNet and SLEAP implemented on U-Net. (E) Accuracy of real-time estimation of head position (top) and tailbase position (bottom) of the mouse during Barnes Maze behavior for each of the three models. (F) Accuracy of real-time estimation of head position (top) and tailbase position (bottom) of the mouse during open field behavior for each of the three models. (G) Distribution of position inference time during APA behavior for the three models evaluated. (H) Distribution of position inference time during Barnes Maze behavior for the three models evaluated. (I) Distribution of position inference time during open field behavior for the three models evaluated.

To evaluate the accuracy of three models - DLC implemented in MobileNetV2, DLC implemented in ResNet, and SLEAP implemented on U-Net, we compared estimation head position (**Figure 2D**) and estimated base of tail position (**Figure 2E**) to 50 randomly selected manually annotated images in each of the three behavioral assays.

Of the three models, DLC implemented in MobileNetV2 consistently demonstrated superior performance across all tasks. In the APA task, DLC implemented in MobileNetV2 achieved perfect accuracy (100% precision and sensitivity) for both head and tailbase predictions. Similarly, in the Barnes Maze task, DLC implemented in MobileNetV2 achieved 100% accuracy for head prediction, and 97.87% and 95.83% precision and sensitivity, respectively, for tailbase prediction. In the open field, DLC implemented in MobileNetV2 again achieved 100% accuracy for both head and tailbase predictions.

We also determined the time taken for estimation or inference time. Mean inference time for detecting head and tail position in an image frame was 61.3 ± 50.5 ms for DLC implemented in MobileNetV2. In comparison, DLC implemented in ResNet took 155.7 ± 135.9 ms, and SLEAP implemented in U-Net took 91.0 ± 48.2 ms. Based on these results, we concluded that the DLC implementation in MobileNetV2 model was ideal for real-time computer vision feedback for the motorized commutator.

### CV feedback allow active compensation of commutator position and rotation angle in response to mouse movement and heading direction

We next evaluated the ability of the CV-guided motorized commutator to actively compensate and adjust the position and angular orientation of the commutator in response to movement of mice in the behavioral arena.

To assess the combined translation and rotation capabilities, mice were tracked along a long linear maze measuring 1.2 m long (**Figure 3A, Supplementary Video 2**). The length of the maze was virtually divided into 8 segments, each ∼150 cm long, and the position of the commutator was adjusted when mice moved from one segment to the next. The heading angle of the commutator was also adjusted when mice accumulated heading angle exceeded a threshold.

**Figure 3.**
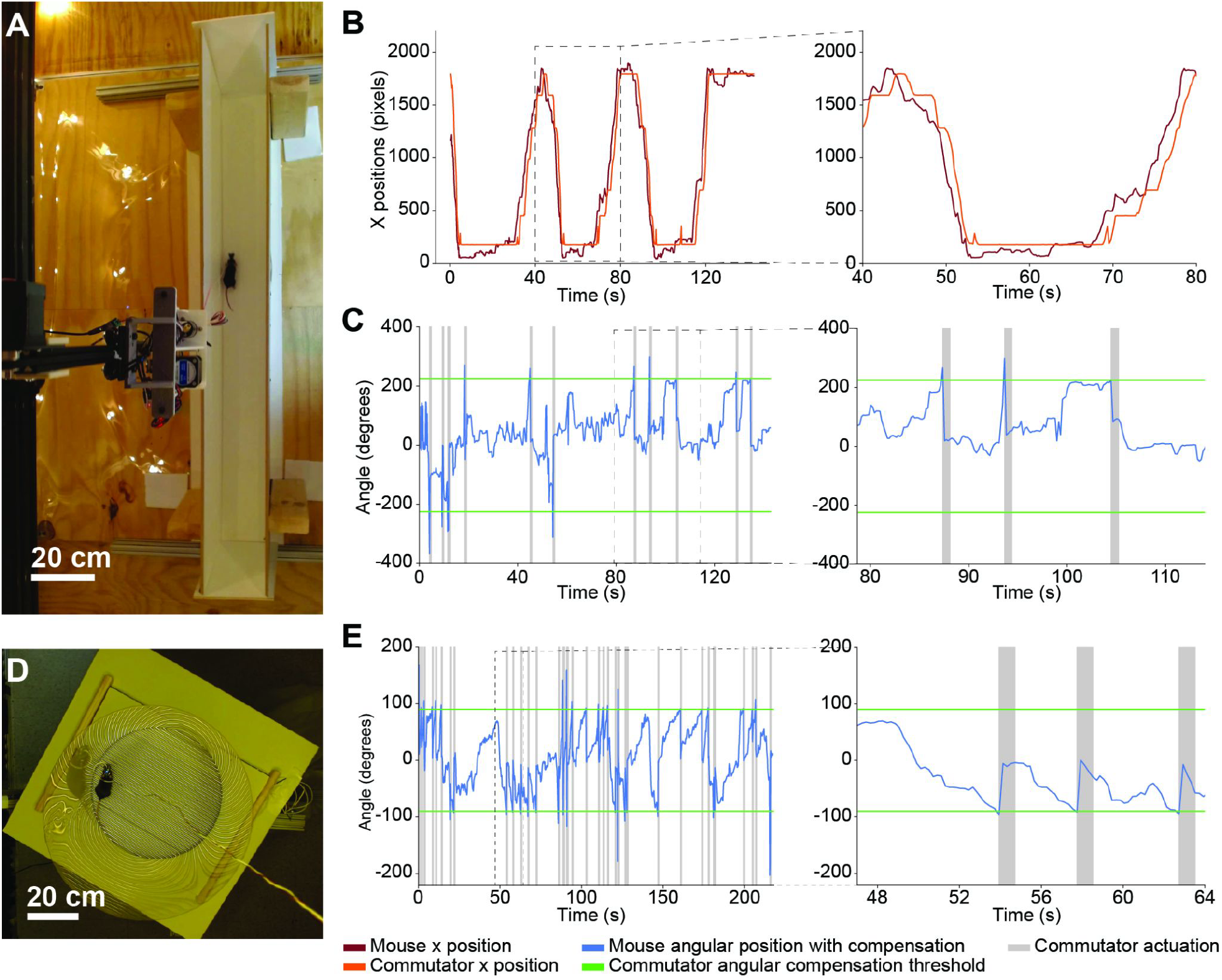
CV-guided active translation and commutation: (A) Still image captured from the overhead video camera of mouse in a 1.2 m long linear track arena. (B) Plot of estimated mouse position in linear track as shown in (a) and the position of the commutator. *Left:* positions over the whole trial, *Right:* highlighting time duration indicated in dashed rectangle in the plot on the left. (C) Plot of estimated mouse heading direction in the linear track arena as shown in (A) and the angular position of the commutator. *Left:* angular positions over the whole trial, *Right:* highlighting time duration indicated in dashed rectangle in the plot on the left. (D) Still image captured from the overhead video camera of mouse in an active place avoidance arena. (E) Plot of estimated mouse heading direction in the active place avoidance arena as shown in (D) and the angular position of the commutator. *Left:* angular positions over the whole trial, *Right:* highlighting time duration indicated in dashed rectangle in the plot on the left.

A plot of the real-time position estimate of the mouse within the linear track and the position of the commutator directly above it is shown in **Figure 3B**. Within the same experiment, the mouse heading angle was also estimated and accumulations of cumulative head motion of 225 degrees resulted in compensation of the commutator to mitigate wire entanglement (**Figure 3C**). We were able to reliably run trials lasting 24 minutes, with no errors in tracking and active compensation observed.

To assess just the rotation capabilities of the CV-guided motorized commutator, we repeated the experiment in the APA task. Mice were tracked in a circular maze of diameter 34.5 cm (**Figure 3D, Supplementary Video 1**). The heading angle of the commutator was adjusted when mice accumulated heading angle exceeded a threshold.

A plot of the real-time heading angle estimates of the mouse within the circular track and the heading angle of the commutator directly above it is shown in **Figure 3E**. Accumulations of cumulative head motion of 90 degrees resulted in compensation of the commutator to mitigate wire entanglement (**Figure 3E**). We were able to reliably run 150 trials ranging from 10-45 minutes, cumulatively 30 hours, with no errors in tracking and active compensation observed. Thus, CV guided active commutation is a reliable method applicable in a wide variety of behavioral assays.

### Stable in vivo imaging during active commutation

We performed wide-field imaging of calcium activities across the whole dorsal cortex using a miniaturized mesoscale imaging device^1^ (**Figure 4A**) while routing the digital interlink through the motorized commutator. We were able to stably record Calcium dynamics at 30 frames per second with only 25 frames dropped during a recording lasting 10 minutes (0.13% of a total of 18,000 frames). Qualitatively, the recorded Calcium activities in the regions of interest distributed throughout the cortex were similar to those acquired with the same device in previous studies^1,31^. We recorded in total 40 minutes, resulting in 130 frame drops (0.18% of 72,000 frames), again comparable to results obtained in our previous work that used a commercial commutator. Note, the frame drop rate is a function of acquisition speed and the configuration of the data acquisition hardware, as well as the efficiency of the computer USB disk drive speed. The frame drops we measured in our motorized commutator system (JP41-119-01HW, Jinpat Electronics) is comparable to the frame drop rate we have observed when using a commercial static commutator system (Carousel Commutator 1x DHST 2x LED, Plexon Inc)^1^. Our modular design can be modified to incorporate other slip rings or commutators with additional data transfer capabilities. Further users will need to pay attention to the data acquisition hardware for efficient data transfer with minimal frame losses.

**Figure 4.**
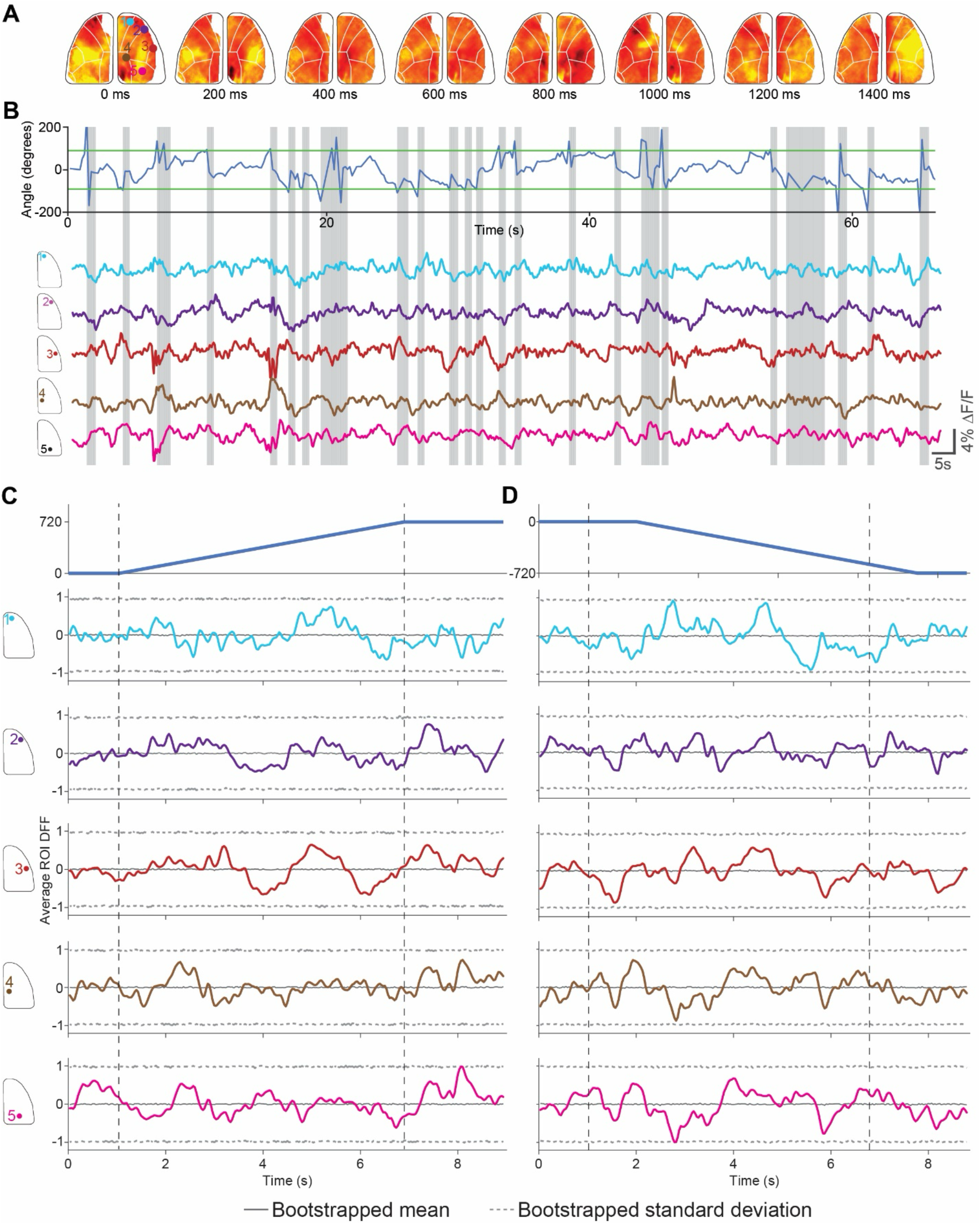
Wide-field calcium imaging in freely behaving mice during active commutation. (A) Pseudo-color DF/F z-score heat maps showing calcium activity progression during an active place avoidance trial during active commutation. (B) *Top:* Mouse body angle tracking during a trial in the active place avoidance task. Gray lines denote active commutation periods to account for mouse angle changes greater than 90 degrees, highlighted in green. *Bottom*: Average DF/F z-score maps plotted for a wide range of regions of interest following the Allen Brain Atlas across one hemisphere of the brain. Gray lines denote active commutation periods to account for mouse angle changes greater than 90 degrees. (C) *Top*: Peri-event time histograms for the average of 10 clockwise rotations of 720 degrees with the active commutator. *Bottom*: Peri-event time histograms for the corresponding average DF/F z-score across 5 regions of interest in the 10 clockwise rotations. Solid color lines indicate the average DF/F z-score for each region of interest. The gray solid line indicates the average of 1000 randomized bootstraps of the DF/F z-score data for the entire trial taken during the commutation period for each region of interest. The gray dashed line indicates the standard deviation of 1000 randomized bootstraps of the DF/F z-score data for the entire trial taken during the commutation period for each region of interest. (D) *Top*: Peri-event time histograms for the average of 10 counterclockwise rotations of 720 degrees with the active commutator. *Bottom*: Peri-event time histograms for the corresponding average DF/F z-score across 5 regions of interest in the 10 counterclockwise rotations. Solid color lines indicate the average DF/F z-score for each region of interest. The gray solid line indicates the average of 1000 randomized bootstraps of the DF/F z-score data for the entire trial taken during the commutation period for each region of interest. The gray dashed line indicates the standard deviation of 1000 randomized bootstraps of the DF/F z-score data for the entire trial taken during the commutation period for each region of interest.

The active commutator turns on the rotational motor when a threshold of 90 degrees of rotation—threshold changes depending on the arena type and size— is exceeded from the starting heading direction angle. Active commutation occurs at a speed of ∼100 deg/s, which we heuristically determined to be slow enough to be imperceptible to the mouse. Nevertheless, the active commutation itself may result in interfering with the neural imaging experiments in two ways - the untwisting of the wired data cable during active compensatory motion may result in mechanical displacement of the imaging device with respect to the skull and the brain. Second, the compensatory motion may be perceptible to the mouse and result in neurophysiological response to the same that might result in artifacts.

Calcium imaging data analysis pipelines typically incorporate motion correction algorithms to account for mechanical displacements of the imaging device with respect to the brain. In our analysis pipeline, we used a rigid body error correction scheme that can correct for lateral displacements of the FOV^34^. We quantified the overall changes in x and y displacement of the FOV as detected by the algorithm throughout the open field behavior trial and specifically around the clockwise (CW) and counterclockwise (CCW) compensatory motion epochs (**Figure 4B**). For the entire trial lasting 10 minutes, average delta X corrections were -1.67 ± 3.3 μm, and average delta Y corrections were 9.04e-4 ± 0.11 μm, respectively. Within the compensatory rotation epochs lasting 140 seconds, average delta X corrections was -2.0 ± 3.2 μm, and delta Y corrections was -1.00e-3 ± 0.06 μm, which is not visually different from the whole trial averages.

We next evaluated Calcium activity during the active compensation epochs. Widefield imaging with the mini-mScope^1^ allows us to look at both global cortex-wide changes, but also at regions of interest located at multiple sensory cortices and motor cortices. All these specific regions of interest may have neurophysiological changes in response to a perception of the compensatory motion. **Figure 4C** and **Figure 4D** show peri-event calcium activity histograms of multiple selected ROIs during CW and CCW active compensation epoch. We found that at a frame-by-frame time scale, average calcium activity with not significantly different (Bonferroni correction, 1000 random bootstraps) from whole trial bootstrapped calcium activity traced for all the ROIs analyzed. Thus, we can conclude that active compensation does not introduce any neurophysiological artifacts.

## Discussion

We present a CV guided active translating commutator that can track and move along with a mouse in behavioral arenas that are large (> 1 m), while minimizing cable length, and using overhead behavioral cameras that simultaneously track the behavior of the animal. The commutator was tested extensively in three separate behavioral experiments - where mice were tracked over large arenas (Barnes maze), in standard open field environments where most of the of the natural repertoire of behaviors were recapitulated, and the active place avoidance tasks where mice performing rapid movements were reliably tracked in real-time over a shocking grid background. We found that the model using the DLC algorithm implemented in MobileNetV2 performed reliably, with 100% precision and sensitivity in the APA task, 98.94% precision and 97.12% sensitivity in the Barnes Maze task, and 100% precision and sensitivity in the open field. Importantly, the model had an inference time of 61.3 ± 50.5 ms when executed on a standard desktop PC. Further, data acquisition using miniaturized microscopes was performed with the same computer. Thus, this is an approach that can be implemented in a wide variety of behavioral experiments using minimal computational resources. In total, we show that active commutation can be achieved without errors in 24 minutes of testing in a linear track, and over 30 hours testing in the APA task.

The CV-guided active commutator uses inexpensive open-source hardware elements for translation and rotation of the slip ring. The linear actuator and the rotary actuators both use the OpenBuilds stepper motors and gantries, which are modular, and can be adapted for much larger arenas. While we implemented translation in one direction in this study, in the future, the approach can be extended to translation in 2 directions. Thus, miniaturized devices with short length (< 1m) wired tethered can be used to interface with head borne devices when mice are exploring significantly larger arenas. In comparison to existing active commutation approaches, our approach simplifies the experiment, but does not require any modifications to the head borne imaging and recording devices. Thus, the approach is compatible with most head borne devices that are already being used by neuroscience laboratories.

While we strived to test the CV guided active commutator in several behavioral contexts, it is possible implementing this in behavioral assays where mice need to be tracked in arenas offering low contrast may result in lesser accuracy and precision in estimating position and heading direction. We incorporated a manual override option to mitigate this issue. Second, the view of the overhead camera can be occluded by the wired tethers occasionally. This may potentially be mitigated by imaging the mice from the bottom of the arena if compatible with the behavioral test.

## AUTHOR CONTRIBUTIONS

IO, SBK conceptualized the technologies. IO, DS developed the technology, IO, DS, KS and ML performed experimental testing. OI, DS, KS, ML, SBK wrote the manuscript.

## SOFTWARE AND CODE AVAILABILITY

All CAD files, code and software associated with this manuscript are available at https://github.com/bsbrl/Motorized-Commutator.git

---

**Figure S1.**
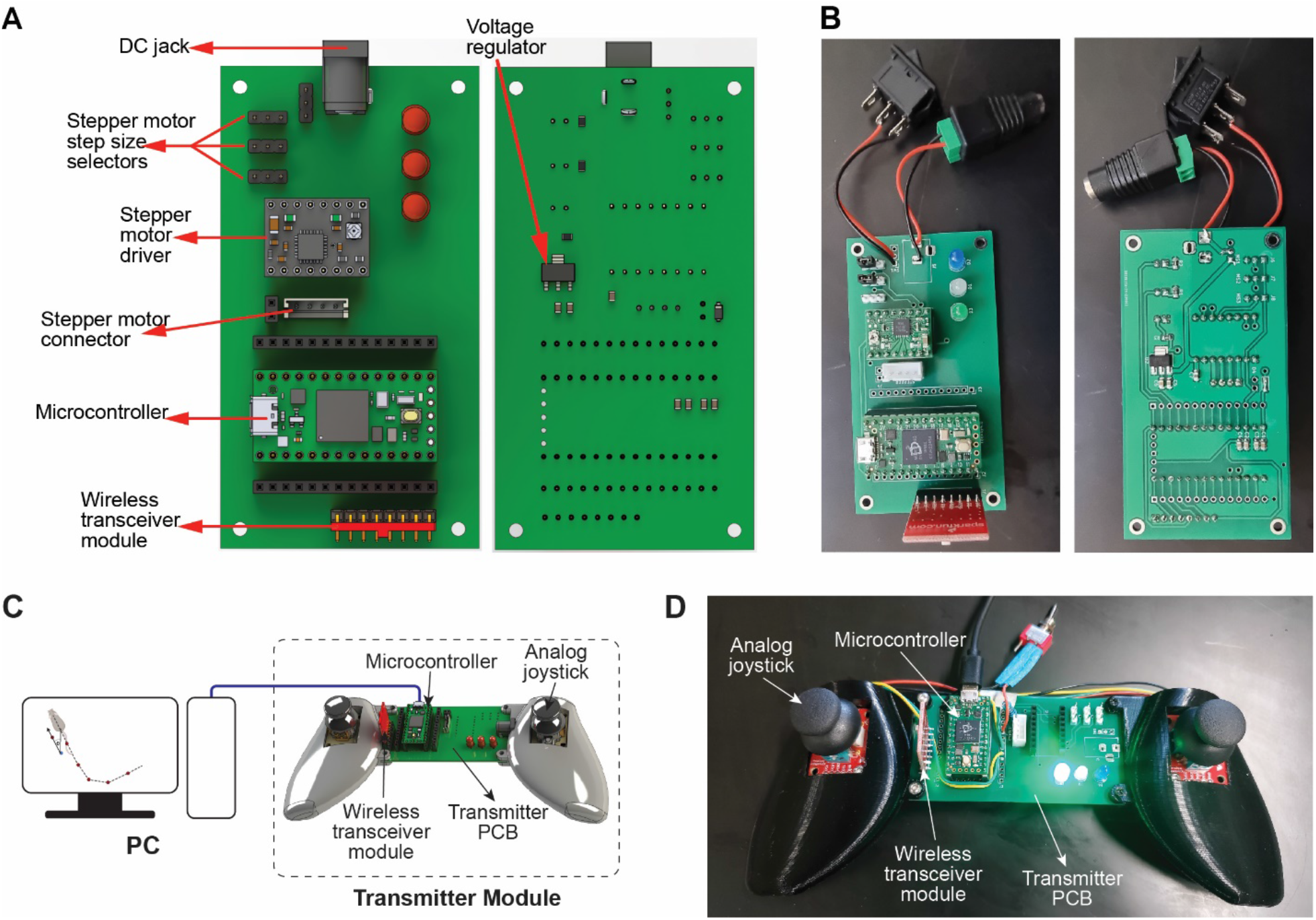
Active motorized commutator PCB. (A) Detailed top and bottom view of the motorized commutator PCB. (B) Top and bottom photo of a fully assembled motorized commutator PCB. (A) This panel illustrates the schematic connections between the personal computer (PC) and the transmitter module within the system. In the active operation mode, the transmitter module receives movement commands from the computer vision tracking algorithm and transmits these commands wirelessly to the motorized commutator and/or gantry in the experiment arena. In the passive operation mode, the transmitter module only gets power from the PC and control of the motorized commutator is done through the joysticks. Mode transition is enabled by the actuation of a toggle switch. (B) Photo of the transmitter module.

**Figure S2.**
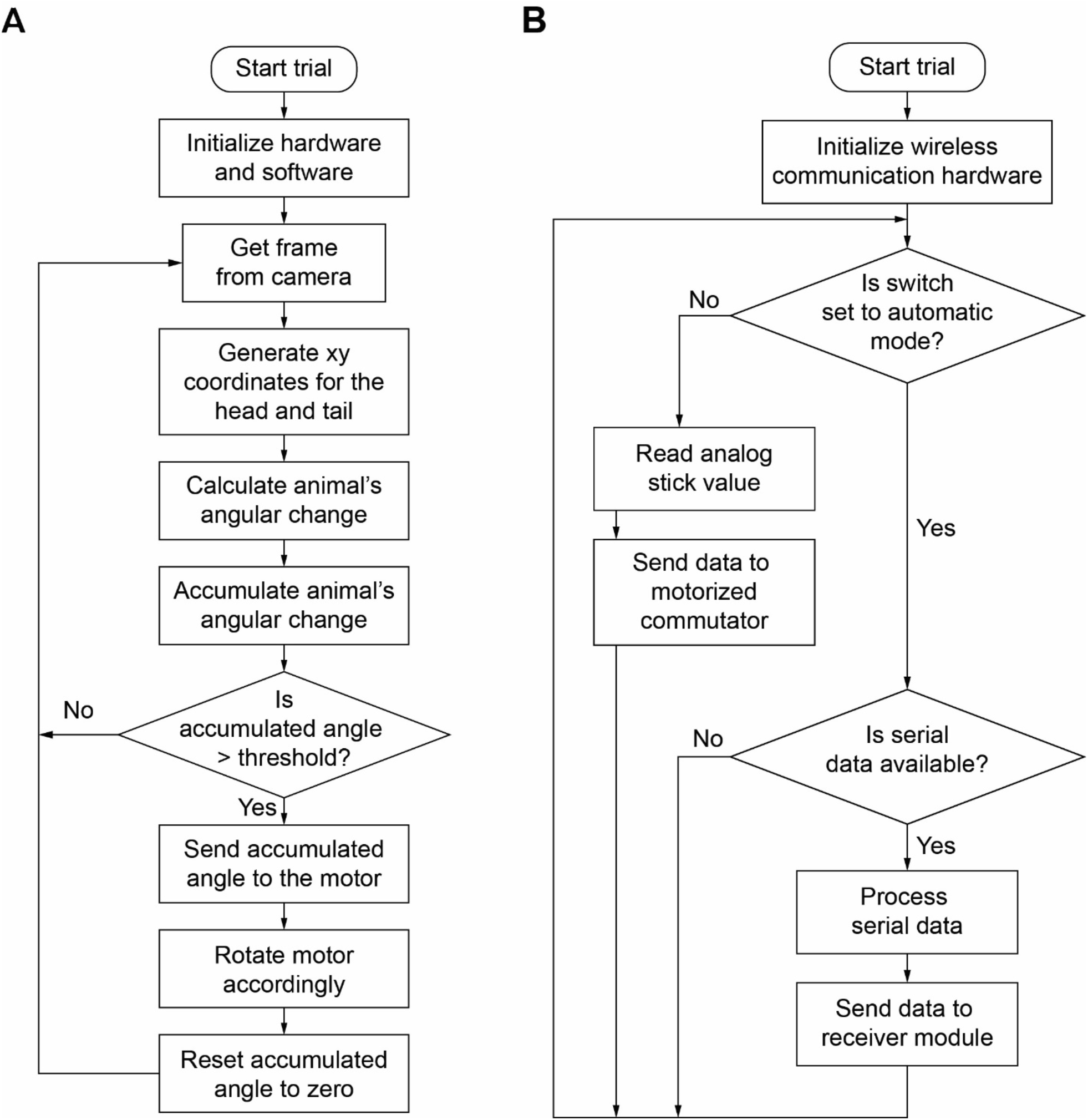
Schematic representation of the cv guided active commutator control algorithm. (A) System flow chart showing how the cv guided active commutator algorithm works. (c) System flow chart showing how the software on the transmitter module interacts with the cv guided active commutator algorithm and the commutator hardware.

